# Genome-wide CRISPRi Screen in Human iNeurons to Identify Novel Focal Cortical Dysplasia Genes

**DOI:** 10.1101/2023.12.13.571474

**Authors:** Andrew M. Tidball, Jinghui Luo, J. Clayton Walker, Taylor N. Takla, Gemma L. Carvill, Jack M. Parent

**Author notes:** Corresponding authors: Andrew M. Tidball, PhD and Jack M. Parent, MD. **Email:** or. **Author Contributions:** A.M.T. and J.M.P. designed research; A.M.T., J.L., J.C.W., and T.N.T. performed research; A.M.T., J.L., and G.L.C. analyzed the data; and A.M.T. wrote the paper with edits from J.M.P. **Competing Interest Statement:** The authors declare no conflict of interest. The funders had no role in the design of the study; in the collection, analyses, or interpretation of data; in the writing of the manuscript; or in the decision to publish the results.

## Abstract

Focal cortical dysplasia (FCD) is a common cause of focal epilepsy that typically results from brain mosaic mutations in the mTOR cell signaling pathway. To identify new FCD genes, we developed an *in vitro* CRISPRi screen in human neurons and used FACS enrichment based on the FCD biomarker, phosphorylated S6 ribosomal protein (pS6). Using whole-genome (110,000 gRNAs) and candidate (129 gRNAs) libraries, we discovered 12 new genes that significantly increase pS6 levels. Interestingly, positive hits were enriched for brain-specific genes, highlighting the effectiveness of using human iPSC-derived induced neurons (iNeurons) in our screen. We investigated the signaling pathways of six candidate genes: *LRRC4*, *EIF3A, TSN, HIP1, PIK3R3,* and *URI1*. All six genes increased phosphorylation of S6. However, only two genes, *PIK3R3* and *HIP1,* caused hyperphosphorylation more proximally in the AKT/mTOR/S6 signaling pathway. Importantly, these two genes have recently been found independently to be mutated in resected brain tissue from FCD patients, supporting the predictive validity of our screen. Knocking down each of the other four genes (*LRRC4, EIF3A, TSN, and URI1*) in iNeurons caused them to become resistant to the loss of growth factor signaling; without growth factor stimulation, pS6 levels were comparable to growth factor stimulated controls. Our data markedly expand the set of genes that are likely to regulate mTOR pathway signaling in neurons and provide additional targets for identifying somatic gene variants that cause FCD.

**Significance Statement:** Focal cortical dysplasia (FCD) is a common cause of focal epilepsy due to somatic variants in mTOR pathway genes in FCD brain tissue. Unbiased sequencing to identify novel FCD genes is challenging since these variants are often in a small subset of neurons. To overcome this challenge, we used an *in vitro* genetic screen in human neurons using an FCD biomarker, uncovering genes that increase neuronal mTOR signaling when their expression is lost. Two candidate genes were mutated in patient FCD brain tissues in a recent study, supporting a causative role for these genes in FCD. Our work suggests gene candidates for somatic variant analysis in FCD tissue and indicates added value for genetic screening in neurons over cell lines.

## Introduction

Focal epilepsies comprise the majority of epilepsy cases (1). Epilepsy in 5-25% of these patients is caused by focal cortical dysplasia (FCD), a localized malformation of cortical development that is often resistant to anti-seizure medications (2). Unlike germline epilepsy-related pathogenic variants that can be identified by sequencing blood samples, FCD is primarily caused by somatic variants, which occur only within the cortical malformation (3, 4). According to current estimates, putatively causative variants are identified in only ∼30% of FCD1 and ∼60% of FCD2 cases (5). Therefore, some of the remaining cases likely have mutations in unknown FCD genes. In patient surgical specimens, FCD-causing variants are only found in a small percentage of cells and rarely in peripheral tissues such as blood. The variant allelic fraction for these pathogenic variants can range from 0.3-18% in bulk tissue but is closer to 50% in neurons harboring FCD pathology (4, 5). Consequently, identifying novel causative genes is extremely challenging due to the limited availability of brain tissue and the need for high read-depth. One must also distinguish pathogenic from benign somatic variants that abound in all tissues, including brain, or rely on laser capture microdissection to sequence DNA solely from cortical neurons that are identified as abnormal (6). For these reasons, causative variant detection in FCD tissue typically relies on screening for mutations in known FCD genes. Therefore, novel approaches are needed to address the question: what genes are we missing in human FCDs?

To overcome these challenges, some recent studies have utilized high read depth and large patient tissue banks to validate new FCD genes (7, 8). As a complementary alternative strategy, in this study we developed an *in vitro* forward genetic screen to identify novel potential FCD-related genes. Most known FCD-related genes are negative regulators of mTOR signaling (except for hyperactivating missense mutations in positive regulator genes like *MTOR* and *PIK3CA*); thus, we sought to identify genes where loss of function (LoF) would lead to mTOR hyperactivation as measured by phosphorylation of the downstream target ribosomal protein S6, a biomarker for FCD2 tissue (9, 10). This strategy has been used in previous screens for mTOR regulators in yeast, human tumors and immortalized cells (11–13). However, unlike these previous studies, we preformed our screen in a more functionally relevant system: human neurons, by utilizing transcription factor-induced neuron (iNeuron) methods (14, 15) and CRISPR interference (CRISPRi). CRISPRi screening in iNeurons was first accomplished by the Kampmann laboratory in 2019 as the initial example of CRISPRi screening performed in a differentiated, human cell type derived from induced pluripotent stem cells (iPSCs). The screen identified genes essential for neuronal survival (16). Unlike other mTOR screens, we performed our screening without amino acid or glucose depletion since FCD causes upregulation of S6 phosphorylation in neurons under normal nutrient conditions. We also focused on negative regulators of mTOR activity rather than positive regulators, which were the focus of several previous studies (11, 12). This LoF approach should be optimal in our model system because neurons have low mTOR activity under basal conditions compared with high basal activity in immortalized cells, and CRISPRi can reduce gene expression of on-target genes by 90-99% without inducing gene mutations and with minimal off-target effects (17).

Using whole genome and candidate gRNA library screens, we successfully identified several known FCD-related genes and novel candidates that upregulated pS6. We further characterized 6 of the novel genes by the In-Cell Western Assay and immunoblotting assays to identify other affected phosphoproteins in the AKT/mTOR/pS6 pathway for each gene knockdown. Two of our characterized genes were found to affect AKT phosphorylation and variants in both genes have been identified in FCD patient brain tissue in a recent study (7). The other 4 genes were found to make the iNeurons resistant to the loss of GDNF-dependent S6 phosphorylation, through an AKT and ERK-independent mechanism.

## Results

### CRISPRi iNeuron iPSC line generation

In this study, our goal was to identify potential novel regulators of mTOR signaling in young neurons as candidate FCD-related genes. We took advantage of established FCD-related genes for internal validation of our screening approach. We anticipated that the screen would provide evidence for genes with variants in a limited number of patients and unveil novel candidate FCD-related genes that can be subjected to deep sequencing in patient samples. To ensure reliability and accuracy of our screening, we designed a human neuronal system that tightly controls gene expression. To achieve this, we used CRISPRi technology, which allowed for efficient knockdown of gene expression from both alleles, while minimizing off-target effects and toxicity associated with traditional CRISPR-Cas9 screens that involve double-stranded DNA breaks. An overview of our screening strategy is schematized in **Figure 1A**. Specifically, we used the KRAB-dCAS9 fusion protein, originally developed by the Weissman lab (18), and constructed a targeting vector for the human *AAVS1* safe-harbor locus with KRAB-dCas9 under the control of the tetracycline-responsive element (TRE) promoter (**Figure S1A**). We transfected human foreskin fibroblasts with this vector and TALENs for the *AAVS1* insertion site, and then selected cells that incorporated the cassette using puromycin selection. We confirmed insertion in the correct locus by PCR. To establish a human neuronal system, we generated doxycycline-inducible neurons (iNeurons) from human induced pluripotent stem cells (iPSCs) (15). In these cells, a doxycycline (dox)-inducible Tet-On system drives the expression of *Ngn1/2* yielding a relatively homogeneous population of excitatory cortical-like neurons (15). We transfected the KRAB-dCas9 fibroblasts with pUCM-CLYBL-*Ngn1/2* and TALEN plasmids targeting the *CLYBL* safe-harbor-like insertion site, as shown in the construct map in **Figure S1B**. Cells were selected for mCherry expression using fluorescence-activated cell sorting (FACS), and insertion was confirmed by PCR. Lastly, we reprogrammed these cells into iPSCs by transfecting them with the three pCXLE episomal reprogramming plasmids (19). Thus, dox treatment of the iPSCs both induces neuronal differentiation and activates CRISPRi via KRAB-dCas9.

**Figure 1.**
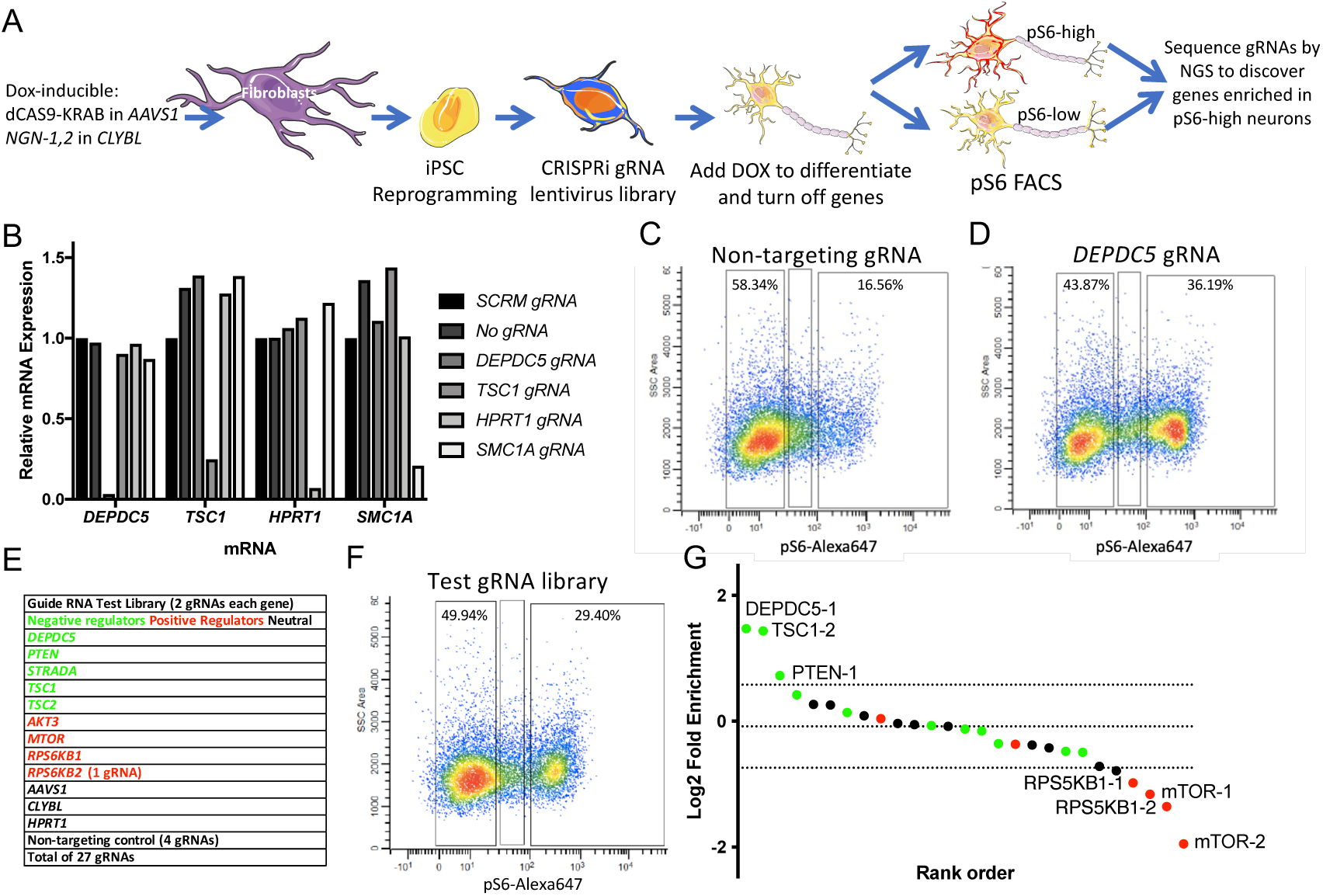
CRISPRi gRNA library screening identifies positive and negative regulators of mTOR signaling in human iPSC-derived iNeurons. (A) Schematic depicting the screening platform design, including the cell line generation, gRNA lentiviral transduction, dox treatment, pS6-based FACS, and next-gen sequencing of the gRNAs. (B) Quantitative RT-PCR data for 4 genes (*DEPDC5, TSC1, HPRT1,* and *SMC1A*) in iNeurons transduced with 5 different gRNAs (NTC, *DEPDC5, TSC1, HPRT1,* and *SMC1A*). (C,D) FACS analysis for pS6-Alexa-647 immunostaining and side-scatter area in iNeurons with non-targeting gRNA (C) or *DEPDC5* gRNA (D). (E) Table of gRNAs in test library comprised of negative regulators of mTOR signaling (green), positive regulators (red), and neutral gRNAs that should have no effect (black). (F) FACS analysis for iNeurons transduced with the test library. The left gate was collected as the pS6-low population (49.9% of total), and the right gate was collected as the pS6-high population (29.4% of total). (G) Log2 fold enrichment of the gRNA sequencing reads in the pS6-high sample vs. pS6-low sample. Dotted lines are mean and 2 SD for the NTC gRNAs. Color coding is the same as for the table in panel (E).

### KN-5 line validation

Clonal lines were selected, and the correct cassette insertions were confirmed by PCR. DNA was also sent for a SNP microarray to identify genomic abnormalities. The KN-5 line was chosen for optimal iPSC growth, lack of genomic abnormalities, and efficient generation of iNeurons after dox exposure (**Figure S1C**). Additionally, qRT-PCR was used to assess whether KRAB-dCAS9 was expressed after dox exposure (**Figure S1D**). While a low level of KRAB-dCAS9 mRNA was expressed without dox exposure, 1 µg/mL dox caused a maximal 20,000-fold increase in expression which was ∼2-fold greater than in the 10 µg/mL exposed cells (**Figure S1D**). Next, to confirm the efficacy of the KRAB-dCas9, we cloned several gRNAs (non-targeting control [NTC], *DEPDC5, TSC1, HPRT1,* and *SMC1A*) from a computationally designed library (20) into the sgRNA expressing plasmid. *DEPDC5* and *TSC1* are mTOR pathway genes, *HPRT1* is an unrelated metabolic gene, and *SMC1A* is a non-FCD-related epilepsy gene. Separate lentiviral vectors carrying each of the gRNA plasmids were generated, and the KN-5 line was transduced with each. Transduced cells were enriched by FACS using blue fluorescent protein (TagBFP) expression on the sgRNA cassette and constitutively expressed mCherry from the cassette in the *CLYBL* locus (**Figure S1E**). The sorted cells were grown until day 7, and RNA was isolated for qRT-PCR. A drastic reduction in mRNA expression was seen for each “on-target” sample, without any off-target effects on the other genes (**Figure 1B**). We also performed immunoblotting for SMC1A protein after CRISPRi in iNeurons transduced with *DEPDC5* or *SMC1A* gRNAs, along with an untransduced control. Only *SMC1A* gRNA-expressing cells showed a large decrease in SMC1A protein expression (86%) (**Figure S1F**) that corresponded nearly identically with the 85% reduction seen in *SMC1A* mRNA (**Figure 1B** and **Figure S1G**).

### iNeurons with constitutive or CRISPRi-induced loss of *DEPDC5* expression have increased S6 phosphorylation

To ensure that loss of expression of mTOR negative regulators would indeed result in elevated mTOR signaling in iNeurons, we generated CRISPR-induced frameshift indels in the *DEPDC5* gene using a combined iPSC reprogramming and gene-editing technique we have previously published (21, 22) applied to KRAB-dCAS9 and *Ngn1,2* cassette integrated foreskin fibroblasts. *DEPDC5* is the most common known cause of FCD and focal epilepsy and is a negative regulator of mTOR signaling in response to the concentrations of specific amino acids (23, 24). The resulting clonal lines underwent PCR for integration of the insertion cassettes and *DEPDC5* genotype. One control line and one homozygous 20 bp deletion line were used to generate iNeurons by dox exposure (**Figure S2A**). After nine days of culture in dox containing medium, the iNeurons were used for western blot analysis (**Figure S2B**). The *DEPDC5-/-* line had a 5-fold increase in pS6 expression compared with the *DEPDC5*+/+ isogenic control line. Cells exposed to improperly buffered HBSS at 5% CO_2_ for 2 hours resulted in a complete loss of S6 phosphorylation.

To assess whether CRISPRi could also lead to a similar increase in S6 phosphorylation, we generated purified iNeurons transduced with NTC, *DEPDC5*, or *SMC1A* gRNA lentivirus. These cells were enriched on day 3 using BFP FACS and used for in-cell western (ICW) quantification for pS6 on a thin bottom 96-well plate after 17 days in culture. The *DEPDC5* gRNA expressing iNeurons showed a 2-fold increase in pS6 expression compared with either the NTC or *SMC1A* gRNAs (**Figure S2C**). It should be noted that we find the ICW can have higher background signal than traditional immunoblotting resulting in reduced estimation of fold-increase. Regardless, this assay was very sensitive in detecting wells with increased pS6 levels. These data indicate that CRISPRi knockdown of an FCD gene results in up-regulation of the FCD biomarker, pS6.

### Known mTOR regulator library screen

Before testing a whole genome gRNA library, we wanted to confirm our ability to screen a multiplex library of gRNAs. To this end, we generated a pooled lentiviral library of 27 individually cloned gRNAs (**Table S1**) targeting known negative and positive regulators of mTOR signaling, as well as “neutral” gRNAs expected to have no effect (**Figure 1E**). Transduction efficiency of the library was assessed using serial dilution and FACS for BFP fluorescence (carried on the gRNA vectors) to achieve an MOI of <0.3 to reduce the probability of cells expressing multiple gRNAs (**Figure S3A,B**). KN-5 iPSCs were transduced with lentiviruses expressing gRNAs for this library, NTCs (negative control), or *DEPDC5* (positive control). After 3 days of dox exposure in iPSC culture conditions, the cells were dissociated, enriched by FACS for BFP fluorescence, and replated into neuronal culture conditions (see Methods). After 4 additional days of culture, all cells achieved a neuronal morphology with robust neurite outgrowth (**Figure S1C**). Cells were again dissociated, fixed, and intracellularly labelled with a primary antibody for pS6 (S235/S236) conjugated to an Alexa-Fluor-647. The cells underwent FACS for Alexa-647 signal. While the NTC gRNA expressing cells had low pS6 signal, we found that many *DEPDC5* gRNA-expressing cells had a dramatic rightward shift in fluorescence intensity (**Figure 1C,D)** with a 20% increase in the pS6-high gate. The identified gates for the pS6-high and pS6-low populations were then used to sort the test gRNA library expressing iNeurons, which also had an increase in the pS6-high population compared with the NTC gRNA iNeurons (**Figure 1E,F**).

After sorting, genomic DNA was isolated from the cells and high-fidelity PCR was used to amplify the gRNA region of the lentiviral DNA. The PCR product was sequenced by NGS and a log2 fold change (LFC) between the 2 gated samples (pS6-high vs. pS6-low) was calculated for each gRNA and plotted as a rank order (**Figure 1G**). We found that the most enriched gRNAs in the pS6-high sample after CRISPRi were known negative regulators of mTOR signaling (*DEPDC5, TSC1, PTEN*) while the most enriched gRNAs in the pS6-low samples were known positive regulators of mTOR signaling (*RPS6KB1, mTOR*). The gRNAs expected to have no effect were closer to the mean values of all the samples (*HPRT1,* NTC, etc.). Of note, several gRNAs for known regulators were near the mean value as well. In fact, another *TSC1* gRNA is the closest value to the mean. This is also the gRNA used for the qRT-PCR experiment in Figure 1B, which was only calculated to reduce the mRNA by 75%, which may not be enough to result in mTOR hyperactivation. For this reason, we decided to examine both individual gRNAs and genes (all gRNAs) in the subsequent whole genome library screening (with 5 gRNAs/gene) since each gRNA can have a different knockdown efficiency resulting in differential effects on mTOR signaling between gRNAs for the same gene (e.g., *TSC1*).

### Whole genome and candidate CRISPRi screen for negative regulators of mTOR signaling

We performed 3 independent whole genome library screens with lentivirus made from the gRNA library containing 5 gRNAs for every gene (104,535 gRNAs total) (20). For each experiment, a positive (*DEPDC5*) and negative (NTC) control were used to define the gates for the pS6-high and low population as performed for the test gRNA library (**Figure 1C,D**). For each experiment we started with more than 100 million cells to ensure sufficient coverage of the entire library (1000 cells/gRNA). However, due to cell loss during dissociation, staining, and sorting, each experiment was underpowered for whole genome coverage. Therefore, during analysis we removed data from gRNAs that had less than 5 total reads in the NGS results for the pS6-high and pS6-low samples for each experiment (55,851 gRNAs across 18,906 genes total) (full CRISPR screen gRNA count data can be found in **Dataset 1**). For each individual gRNA, the LFC between the pS6-high and pS6-low samples and means were calculated across the 3 experiments and presented in rank order (**Figure 2A**). Gene ontology analysis was performed on the rank order for the highest-ranking gRNA for each gene using Gorilla (25), and the highest-ranking terms are shown (**Figure 2B**). Additionally, a rank order for each gene was calculated using data from all gRNAs (> 5 gRNA reads total). The mean rank order between the three experiments was calculated and also used for gene ontology (**Figure 2C**). For each method of analysis, gene ontology identified expected terms for our pS6-based screen (TSC1-TSC2 complex, negative regulation of TOR signaling, and GATOR1 complex). These results provide internal validation to our analysis despite being underpowered for full genome coverage. Therefore, we chose to perform screening on a subset of candidate genes (n= 112) from the library in the same manner using the most significant gRNAs that were also highly ranked by individual gene analysis (candidates are labelled green in **Figure 2A, n=129**).

**Figure 2.**
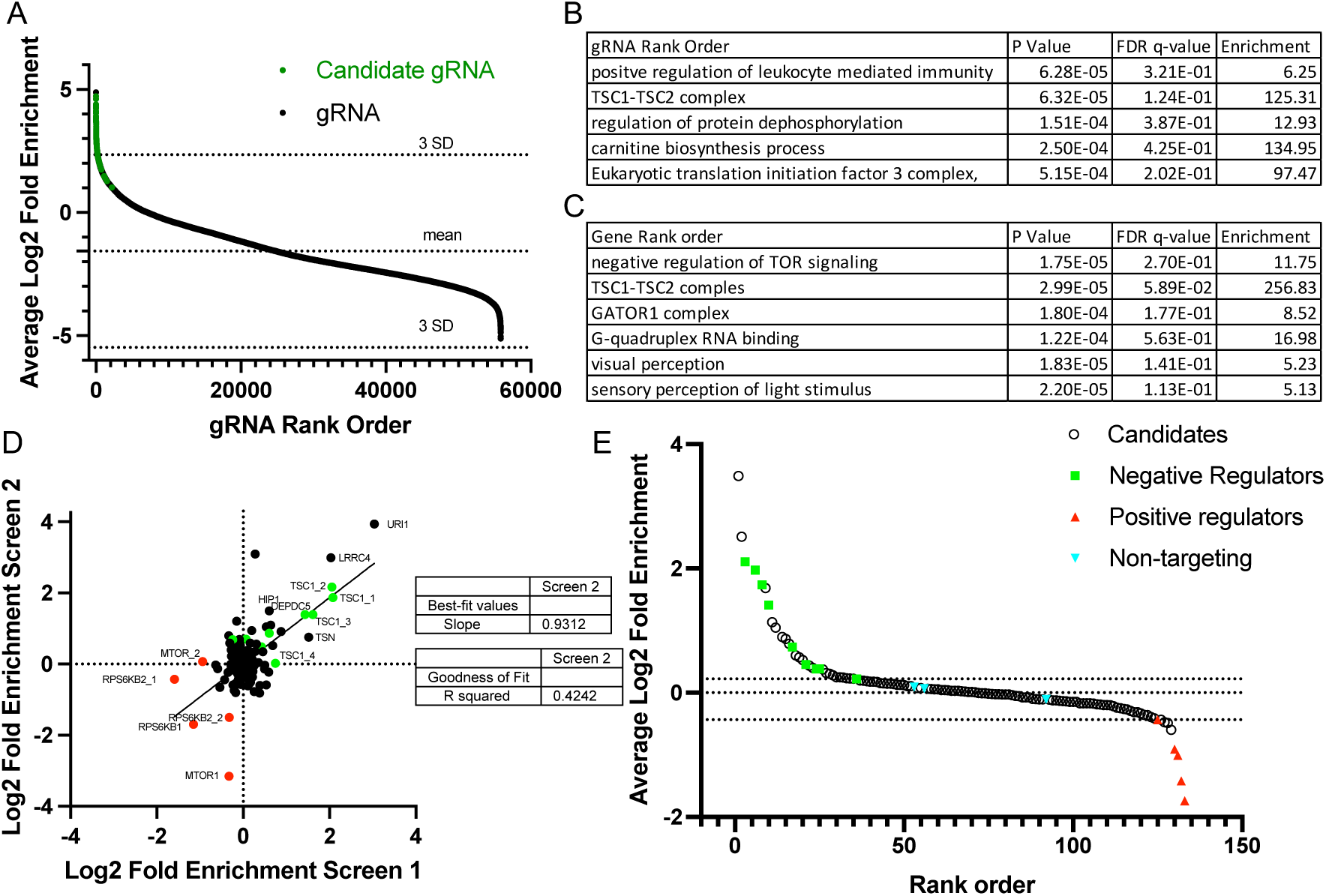
Whole genome gRNA library screening is enriched for known negative regulators of mTOR signaling, and candidate screen identifies gRNAs from novel genes with robust effect on pS6. (A) The average log-2 fold enrichment scores for the whole genome gRNA library. gRNAs with less than 5 total NGS reads combined between the pS6-high and pS6-low samples for all 3 independent experiments were excluded. Candidate gRNAs are labeled in green. (B) Top significant gene ontology terms from the gRNA rank order averaged from the 3 independent experiments. (C) Top significant gene ontology terms from the gene rank order averaged from the 5 gRNAs for each gene across the 3 independent experiments. (D) Plot of log2 fold enrichment in the pS6-high population for the candidate library (129 gRNAs) comparing 2 independent experiments (Screen 1 on x-axis, Screen 2 on y-axis). Green depicts known negative regulators of mTOR signaling, and red depicts known positive regulators of mTOR signaling. (E) Average log2 fold enrichment for the candidate library screen across 3 independent experiments is plotted against rank order in the x-axis.

A library of 129 candidate gRNAs was generated by individual gRNA cloning into the pBA904 CRISPR guide vector that contains a cs1 capture sequence that can be used for features barcode analysis on the 10x Genomics platform (26) (**Dataset 1**). The candidate library contained 112 novel candidate genes and 14 known mTOR regulator gRNAs (targeting *DEPDC5, PTEN, TSC1, TSC2, NPRL2 RPS6KB1, RPS6KB2, mTOR*) and 3 NTC gRNAs. Two individual pS6 screens were performed with NGS identifying 129 gRNAs from each experiment. In addition to NTC and *DEPDC5* gRNA controls, we also included an *MTOR* gRNA, isotype Alexa-647 control antibody, and rapamycin treatment. The isotype control, *MTOR* gRNA, and rapamycin (20 nM for 2 hours) all resulted in a near complete loss of the pS6 high population, while *DEPDC5* and the candidate gRNA library both increased the pS6 high population compared with the NTC gRNAs (**Figure S4**). The LFC for each gRNA is plotted for the two experiments with a linear regression having a R^2^ value of 0.4242 and a slope of 0.9312 indicating the data from the two experiments were consistent (**Figure 2D**).

### Top candidate negative regulators confirmed by in-cell western and immunoblots

The top 19 gRNAs including the top 14 candidate gRNAs (14 genes) from the candidate screening and 5 gRNAs for known FCD-related genes highly ranked in our screen (*PTEN, DEPDC5, NPRL2, TSC1,* and *TSC2*) were used to generate individual gRNA lentiviral vectors. Transduced iNeurons were plated at high density in a 96-well plate and In-Cell Western blot (ICW) was performed using the anti-pS6 (S235/236) antibody. Of the candidates, 8 had significantly increased pS6 compared with NTC (**Figure 3A**). Surprisingly, none of the known FCD-related genes were significantly increased compared to NTC despite the results from the screen. One possible reason is that the iNeurons are exposed to low nutrient levels during dissociation prior to fixation for the FACS-based screen; however, during ICW the iNeurons went directly from nutrient rich media into fixative. Therefore, pS6 levels were likely suppressed low nutrient conditions in control cells while the levels in cells with knockdown of FCD genes were elevated.

**Figure 3.**
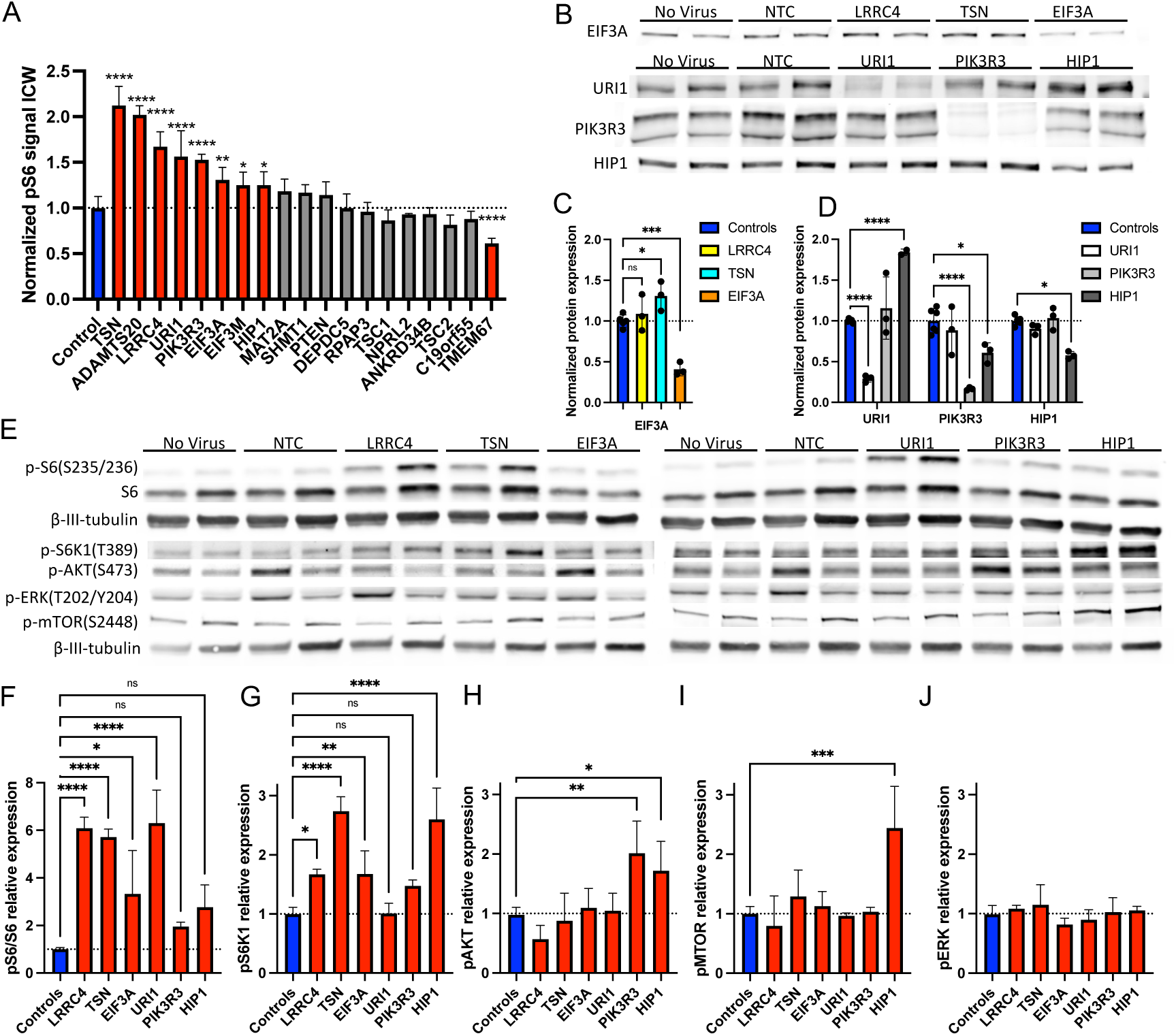
Six candidate genes validated for CRISPRi knockdown and elevated pS6, and the finding of altered upstream AKT/mTOR/S6 signaling in *PIK3R3* and *HIP1* knockdown cells. (A) In-cell western data for 20 gRNAs with n = 4 except for PTEN where n – 3 due to one well peeling/lifting. (B) Representative immunoblots for targeted proteins EIF3A, URI1, PIK3R3, and HIP1 from iNeuron lysates after various gRNA transductions (listed at top of blots). (C) Quantification of EIF3A protein level from 4 independent experiments. The gRNAs used are indicated in the legend. Normalized to beta-III-tubulin. No virus and NTC were combined as “controls.” (D) Quantification of URI1, HIP1, and PIK3R3 protein level from 4 independent experiments. The gRNAs used are indicated in the color legend. (E) Phospho-specific mTOR pathway immunoblot images for iNeurons transduced with gRNAs targeting each gene listed at top. (F-J) Quantification of immunoblot data for 3 independent replicates. (F) The level of pS6(S235/S236) was normalized to the level of total S6 protein. (G) Level of pS6K1(T389) normalized to beta-III-tubulin. (H) Level of pAKT(S473) normalized to beta-III-tubulin. (I) Level of pERK(T202/Y204) normalized to beta-III-tubulin. (J) Level of pMTOR(S2448) normalized to beta-III-tubulin. Side-by-side bands with the same labels are biological replicates from independent transduction/differentiation replicates. Statistical analysis performed by one-way ANOVA with Dunnett’s multiple comparison test. Error bars are means with SD. * p < 0.05, ** p < 0.01, *** p < 0.001, **** p < 0.0001.

While ICW was a useful screening assay for 10’s of gRNA candidates, direct western immunoblotting is the gold standard for confirming changes in protein or phospho-protein abundance. Therefore, we sought to confirm 6 of the 8 novel candidate genes (*TSN, LRRC4, URI1, PIK3R3, EIF3A,* and *HIP1*) using traditional immunoblotting techniques. First, for 4 of the genes (*URI1, PIK3R3, EIF3A,* and *HIP1*) that had well-documented antibodies, we immunoblotted for on-target protein expression (**Figure 3B**). Notably, these cultures were not purified for transduced cells by FACS sorting due to the large number of cells needed to perform western blotting. Instead, we used a much higher MOI for the lentivirus since only single gRNAs were used for each sample. In each case, the on-target expression was dramatically reduced compared with the NTC and off-target gRNA samples (59% reduction for *EIF3A*, 71% for *URI1*, 83% for *PIK3R3*, and 42% for *HIP1*) (**Figure 3C-D**). Interestingly, knockdown of *HIP1* causes an 84% increase in URI1 protein expression while knockdown of *TSN* caused a 31% increase in EIF3A protein expression. These findings suggest either shared molecular mechanisms or compensatory changes. We measured *LRRC4* and *TSN* on-target expression reduction by qRT-PCR and found mRNA levels were reduced 90% and 50%, respectively (**Figure S5A,B**). We also found a second, higher molecular weight band for PIK3R3 that was also greatly reduced in the gRNA-treated cells by the same approximate amount (**Figure 3B**). The size fit with a known fusion readthrough protein from an open reading frame upstream of PIK3R3 called *P3R3URF*. To test if this was the readthrough protein, we performed qRT-PCR for *PIK3R3, P3R3URF-PIK3R3,* and *P3R3URF* transcripts. We only found detectable transcript for *PIK3R3* indicating that this second band is possibly a post-translational modification, but our qRT-PCR does show a 70% reduction in *PIK3R3* mRNA with the on-target gRNA (**Figure S5C**).

We next assessed the phosphorylation of ribosomal protein S6 (S235/236) as well as the phosphorylation of S6 kinase1 (T389), AKT (S473), ERK (T202/Y204), and mTOR (S2448), which are all indicative of their kinase activity. For 4 gRNAs, pS6 normalized to S6 protein was increased compared to NTC gRNA and no virus controls (**Figure 3E,F**). The gRNAs for *HIP1* and *PIK3R3* appeared to increase normalized pS6 levels, but they showed no statistically significant differences from controls. All gRNAs but *URI1* elevated S6 kinase phosphorylation (**Figure 3G**). AKT phosphorylation was increased only after *PIK3R3* or *HIP1* knockdown (**Figure 3H**). This finding is intriguing since *PIK3R3* encodes for p55γ, a subunit of the PI3K complex that phosphorylates AKT. Only *HIP1* knockdown increased mTOR phosphorylation, and no gRNAs caused significant changes in ERK phosphorylation (**Figure 3I,J**). Thus, only *PIK3R3* and *HIP1* seem to affect the upstream *PI3K/mTOR/S6* pathway. Since all known FCD genes regulate mTOR directly or upstream in the GATOR1 or AKT pathways, these two genes are the most likely to be “true” FCD genes. We reviewed the literature for probable pathogenic variants in this small subset of candidate genes in FCD patient tissues and found one study describing multiple patients with missense variants in *HIP1* and *PIK3R3* found in screening and validated sequencing in a single patient each (7). One additional study identified an epilepsy patient with a *HIP1* deletion who had an FCD and dysembryoplastic neuroectodermal tumor (DNT) in the excised epileptic tissue (27). These studies corroborate our findings and suggest that *HIP1* and *PIK3R3* are true FCD genes that should be further studied and included on FCD gene panels.

### Candidate genes cause mTOR resistance to loss of GDNF stimulation

Because our In-Cell Western data showed no elevation of S6 phosphorylation with known FCD genes (*DEPDC5, TSC1, TSC2, NPRL2, PTEN*) (**Figure 3A**), we hypothesized the iNeurons were cultured in conditions that were not amenable to activation of mTOR signaling by these genes. Indeed, we observed under basal conditions in our controls that pS6 is exceedingly low, potentially indicating insufficient growth conditions (**Figure 3E**). Given that neurotrophic factors are important for neuronal health and survival, we had previously begun BDNF and GDNF addition to the iNeuron media on day 8, the day after our pS6 experiments (15). Therefore, we tested whether earlier addition of neurotrophic factors changed the mTOR response of iNeurons to loss of the known FCD gene *DEPDC5* or one of our new candidate genes, *LRRC4,* that is also brain-specific (28). By In-cell Western, we only found a clear increase in pS6 after *DEPDC5* CRISPRi when BDNF and GDNF were present in the culture media (**Figure 4A**) while the opposite was seen after *LRRC4* CRISPRi knockdown in iNeurons (**Figure 4B**). We then sought to determine whether a specific neurotrophic/growth factor(s) stimulated iNeuron mTOR activity. We tested 20 ng/mL treatment with BDNF, GDNF, NT3, or FGF2 (each alone) for 24 hours followed by ICW for pS6. Only GDNF exposure caused a statistically significant increase of 78% in pS6 by ICW (**Figure 4C**). We next tested our 6 novel genes for effects on pS6 in the presence of GDNF. In both no gRNA and NTC gRNA conditions we observed similar, nearly two-fold increases in pS6 levels with GDNF exposure, and cells transduced with the *HIP1* gRNA showed a significant increase with GDNF, while the apparent increase for *PIK3R3* was not statistically significant (**Figure 4D**). However, *LRRC4, TSN, EIF3A,* and *URI1* knockdown all showed similarly elevated pS6 with or without GDNF, indicting resistance to the loss of GDNF effects on mTOR signaling (**Figure 4D**).

**Figure 4.**
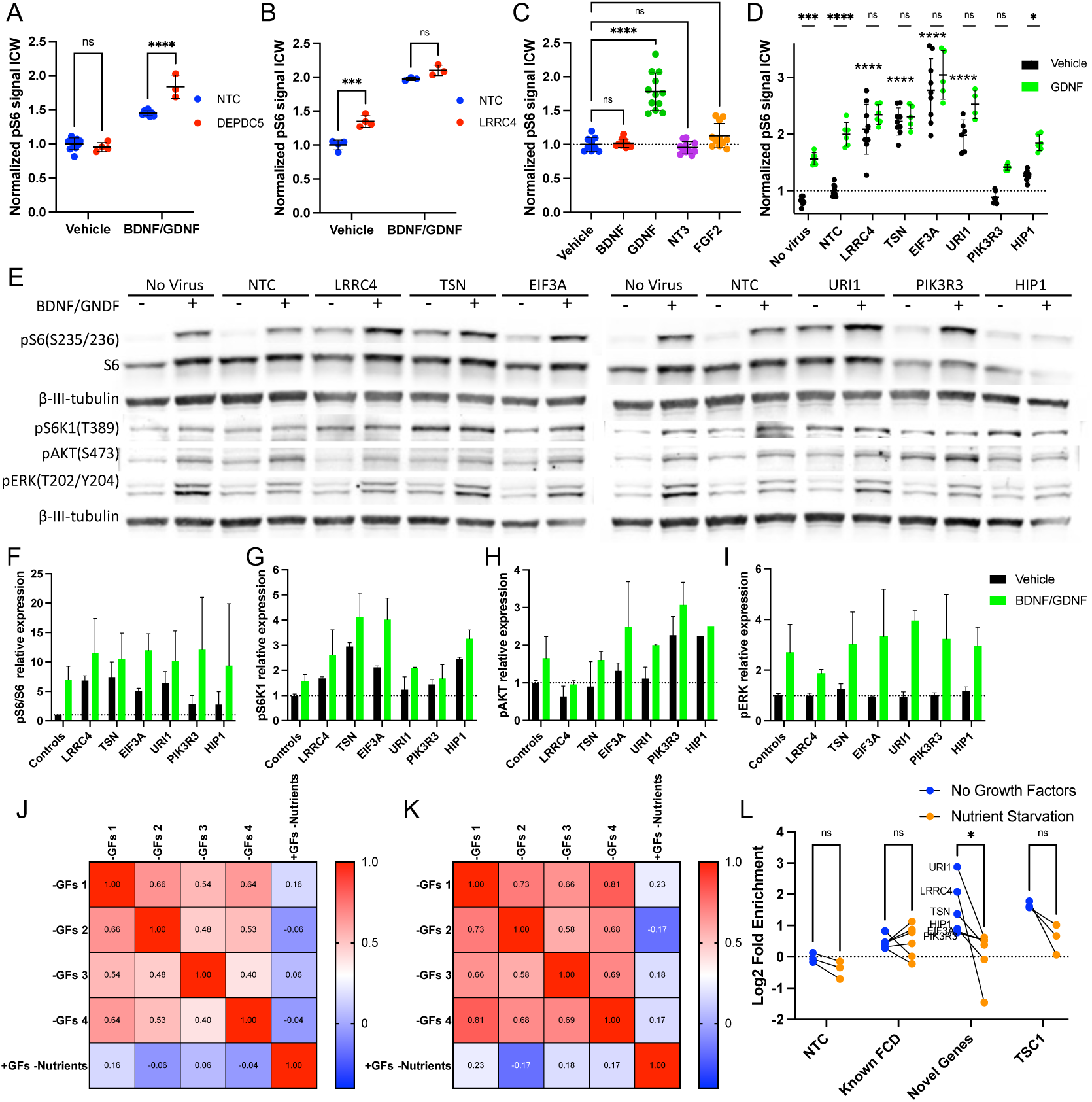
Validated candidate genes cause resistance to loss of glial-derived neurotrophic factor (GDNF) but not nutrient deprivation. (42) ICW for pS6 in iNeurons. Each dot indicates a single well. Each graph is from a single experiment that has been replicated with similar results (not shown). Statistical analysis performed by two-way ANOVA with Tukey’s multiple comparisons test or one-way ANOVA with Dunnet’s multiple comparisons test. Error bars are means with SD. * p < 0.05, ** p < 0.01, *** p < 0.001, **** p < 0.0001. (A) iNeurons were transduced with either NTC or *DEPDC5* gRNA and with or without BDNF and GDNF added to the culture media. (B) iNeurons were transduced with either NTC or *LRRC4* gRNA and cultured in the presence or absence of BDNF and GDNF. (C) iNeurons were treated with BDNF, GDNF, NT3 or FGF2, and only GDNF increased pS6. (D) iNeurons were transduced with the gRNAs labeled on the x-axis and were cultured with (green) or without (black) GDNF. Comparisons in the top row indicate statistical significance for vehicle vs. GDNF conditions for each gRNA, while individual asterisks below indicate significance from the vehicle-treated NTC. (E) Representative immunoblotting for phosphorylated proteins associated with increased mTOR pathway signaling in iNeurons transduced with gRNAs targeting each gene listed at the top, and cultured in the presence (+) or absence (-) of BDNF and GDNF. (F-I) Quantification of immunoblot data for 2 independent replicates. (F) The level of pS6(S235/S236) was normalized to the level of total S6 protein. (G) Level of pS6K1(T389) normalized to beta-III-tubulin. (H) Level of pAKT(S473) normalized to beta-III-tubulin. (I) Level of pERK(T202/Y204) normalized to beta-III-tubulin. (J) Pearson’s correlation matrix for candidate screen experimental data for the 4 experiments without growth factors and the one experiment with growth factors but with nutrient deprivation. (K) The same plot as in (J) but limited to data from validated candidate genes, known FCD genes, and controls. (L) The log2 fold enrichment in the pS6 high population by gRNA with either growth factor or nutrient withdrawal. Statistical analysis performed with multiple Wilcoxon tests (paired non-parametric). Error bars are means with SD. * p < 0.05, ** p < 0.01, *** p < 0.001, **** p < 0.0001.

To confirm this result, we performed traditional western blot analysis to investigate the effects of BDNF/GDNF exposure in the gRNA treated iNeurons. We found a robust 7-fold increase in pS6/S6 for pooled controls exposed to BDNF/GDNF, which further exemplifies the greater sensitivity of immunoblot compared with ICW (**Figure 4E,F**). Exposure to 4 of the gRNAs (*LRRC4, EIF3A, TSN,* and *URI1*) yielded similar levels of pS6 without BDNF/GDNF exposure as the controls *with* BDNF/GDNF exposure, while *PIK3R3* and *HIP1* had elevated pS6/S6 levels without BDNF/GDNF but to a lesser extent. Exposure to BDNF/GDNF led to an ∼ 50% increase above these already elevated levels for all of the genes except *HIP1*; however, it should be noted that some percentage of each culture is not transduced with the gRNA lentivirus and likely has normal pS6 at basal conditions that can be elevated with growth factor exposure. Similar results were seen for phosphorylation of S6 kinase1 (T389) with growth factor stimulated phosphorylation in controls being similar to unstimulated levels for several gRNAs (**Figure 4G**). Phospho-AKT was reduced with *LRRC4* knockdown and elevated with or without growth factor exposure after knockdown of *PIK3R3* or *HIP1* (**Figure 4H**). Interestingly, the growth factors caused a marked elevation in ERK phosphorylation for all gRNAs but with no apparent differences compared with controls (**Figure 4I**).

Lastly, to understand whether the 6 genes we have characterized specifically cause mTOR signaling resistance to loss of growth factors, we performed the candidate library screen again after either growth factor deprivation or a 2-hour nutrient deprivation with BDNF/GDNF present. Pearson’s correlation matrixes for the complete datasets (**Figure 4J**) or the subset with clear condition-specific changes (**Figure 4K**) showed a high degree of correlation between the 3 original datasets from **Figure 2D,E** and the new data without BDNF/GDNF (-GFs 4) with an average correlation coefficient of 0.52 and 0.73, respectively. However, there is almost no correlation between these three experiments and the nutrient deprived experiment with BDNF/GDNF present with average coefficients of 0.03 and 0.10, respectively. A heatmap for the log2 fold enrichment for each gRNA in these two datasets is provided in **Figure S6.** We then compared the log2 fold change for the gRNAs between these two experiments for 4 categories of genes (**Figure 4L**). We compared non-targeting controls and found no difference between the two with the average LFC for each around 0 (p = 0.25). We then looked at all the known FCD genes; however, we noticed that *TSC1* gRNAs had a much greater response during growth factor withdrawal than any other FCD gene gRNAs. Therefore, we removed *TSC1* gRNAs from the group and analyzed them on their own. For the FCD gene gRNA group we see a nearly identical average LFC for each condition (p = 0.999). We then grouped the 6 candidate gene gRNAs we have extensively validated (*LRRC4, TSN, EIF3A,URI1, PIK3R3, HIP1*). All 6 gRNAs had reduced enrichment in the nutrient deprived condition with the average Log2 fold change dropping from 1.47 to 0.086. Using a paired non-parametric t-test, we found a significant difference between conditions for this group but not the other three groupings (p = 0.0313). However, *TSC1* shows a remarkably similar response to growth factor withdrawal compared with the candidate genes and a similar response to the known FCD genes with nutrient starvation. These data indicate that we have identified genes that may be specific negative regulators of growth factor-dependent but not nutrient-dependent mTOR signaling in neurons.

## Discussion

In this study, we developed a human neuronal CRISPRi screening platform that can identify genes that alter a particular cell signaling event, the phosphorylation of ribosomal protein S6. Whole genome gRNA library screening resulted in enrichment for mTOR negative regulators, including the TSC1-TSC2 complex and GATOR1 complex genes. Subsequent candidate gRNA screening removed false positives resulting in a refined list of positive hits. By in-cell western and immunoblot assays, we validated 6 novel genes that significantly elevated S6 phosphorylation when knocked down. Interestingly, we have also shown that these negative regulators cause neurons to have elevated pS6 in the absence of neurotrophic factor stimulation, specifically GDNF.

Previous screens for mTOR regulators have either removed nutrients (amino acids and/or glucose) to identify genes that resulted in mTOR signaling resistance (S6 phosphorylation remaining high) (13, 29), screened for genes that reduced basally elevated S6 phosphorylation in a cancerous cell line (11), or screened for genes that resulted in the inability to reactivate mTOR signaling after nutrient-deprivation and refeeding (12). Our screen did not utilize any designed culture condition but rather anticipated that FCD genes result in elevated pS6 regardless of conditions, as seen in FCD brain tissue. This may have only been possible in our culture system because postmitotic neurons have low pS6 compared with immortalized cells lines used in previous publications (11–13, 29). Interestingly, we recognized that at the early timepoint we used for the screen (7 days of dox exposure), iNeurons are already sensitive to neurotrophic factors. Therefore, our screen may have been biased toward genes that negatively regulate pS6 signaling when neurotrophic factors are absent. A benefit of conducting our screen in human neurons is the potential to identify mTOR regulating genes that are only present in these cells. In fact, 3 of the 13 novel genes - *ADAMTS20, ANKRD34B,* and *LRRC4* - are only found in brain tissue according to RNA sequencing in the human protein atlas database with barely detectible RNA expression in HEK293 cells (28). Other candidate genes from our screen, such as *HIP1*, *TSN* (30), and *PROSER3* (*C19orf55*), show greatest expression in the brain and testes compared with other tissues.

The recent identification of putative pathogenic variants in *HIP1* and *PIK3R3* in patient FCD brain tissue (7) provide validity for our screening to identify novel FCD genes, adds support for the gene variants being causative in FCD, and provides added evidence to investigate the role of these genes in AKT/mTOR/S6 signaling. Somatic and germline variants in a related gene, *PIK3R2*, have been previously associated with the known mTOR hyperactivation-related condition megalencephaly (31). An additional patient reported to have a deletion that includes the *HIP1* gene had a temporal lobe cortical dysplasia found in tissue removed to treat medically refractory focal epilepsy (27). In fact, epilepsy is a common symptom of *HIP1* deletions (32). The HIP1 protein is also known to interact with and regulate the membrane expression of growth factor receptors (33). This regulation may be the reason our growth factor deficient screen identified *HIP1*. These data suggest that the two genes, particularly *HIP1*, play a role in epilepsy etiology, and our study suggests that this is due to *HIP1* negative regulation of mTOR signaling. The healthy control genome database, gnomAD, suggests intolerance for loss of function variants in FCD genes (*DEPDC5, NPRL2, NPRL3, PIK3CA, PTEN, TSC1,* and *TSC2*) as shown by a low average pLI score of 0.30 (range 0.07-0.51), and *HIP1* has a similar score of 0.33. Taken together with the FCD patient sequence data (7), our findings suggest that *PIK3R3* and *HIP1* should be further investigated as FCD genes and potentially added to FCD gene panels.

The other four characterized genes (*LRRC4, EIF3A, TSN, URI1*) likely alter the activity or localization of S6K or RPS6 directly since we did not see an impact on the mTOR pathway except for S6K and RPS6 phosphorylation. In fact, as a part of the eIF3 complex, *EIF3A* is known to interact with both S6K and RPS6, and loss of *EIF3B* or *EIF3C* has previously been shown to increase S6K1 activity (34). In addition, URI1 forms a negative feedback loop for S6K1 activity through its interaction with protein phosphatase 1 gamma (PP1γ). When S6K becomes activated, it directly phosphorylates URI1 causing it to disassociate from PP1γ. PP1γ then desphosphorylates S6K1 causing its inactivation (35). Oddly, *URI1* only has an effect on S6 phosphorylation with no apparent effect on S6K1 phosphorylation in our model system, suggesting some differences from the prior study in HeLa cells. *LRRC4* is a well-known glioma suppressor brain-specific gene that is implicated in the regulation of intracellular signaling, including as a negative regulator of ERK signaling (36). Again, we see different results in our system with no difference in ERK signaling with *LRRC4* knockdown. The fact that some of our candidate genes are specific to the brain and exhibit distinct signaling patterns compared to previous work done in immortalized cell lines underscores the importance of studying signaling regulation in cell types that are relevant to the disease being modeled. By doing so, we can gain a better understanding of how these genes contribute to disease pathogenesis and potentially identify new therapeutic targets.

Our screen led to the discovery of two genes, *HIP1* and *PIK3R3*, that regulate the AKT/mTOR/S6 pathway and have also been found to be mutated in patient FCD tissue. We also found four other novel genes, *LRRC4, TSN, EIF3A,* and *URI1,* that regulate S6 phosphorylation in a new and unique way that may be related to growth factor signaling, although these genes have thus far not been found to be altered in FCD tissue. The CRISPRi iNeuron model provides a platform for exciting future studies with enhanced relevance to FCD and other neurological conditions. For example, the platform and strategy described in this study could be applied to a broader range of signaling events beyond mTOR/S6 activity, including ERK, CREB, AKT, and JAK-STAT pathways. Such work could help identify important genes in these pathways specific to neurons and, thus, be an important tool for cell-type specific functional genomics.

Our list of genes could further be mined for additional useful information. For instance, although not directly addressed here, our screen is bidirectional and can identify genes that reduce mTOR signaling when knocked down. Given our focus on potential FCD genes, the positive mTOR regulators for which knockdown leads to low pS6 were not pursued. However, we note that the most enriched gene in the pS6-low population was *SRSF1*, a gene previously known to activate mTOR (37). SRSF genes have also been identified in a previous mTOR screen (12). *PDPK1* was also identified as the 6^th^ most enriched gene in the pS6-low population and is a known integral part of the PI3K-Akt pathway. Future studies characterizing these genes may be useful to better understand mTOR-related signaling and provide alternative molecular targets for disorders involving the mTOR pathway.

## Materials and Methods

### AAVS1-DOX-KRAB-dCAS9-puro plasmid construction

Puro-Cas9 donor was a gift from Danwei Huangfu (Addgene plasmid # 58409)(38). This plasmid was restriction enzyme digested with AgeI and SalI. The larger band containing the AAVS1 homology arms, tet operator, and puromycin resistance gene were in the larger of the two bands (7184 bp). This band was cut from a 0.7% agarose gel and purified by the Zymo Gel purification kit. pHR-SFFV-KRAB-dCas9-P2A-mCherry was a gift from Jonathan Weissman (Addgene plasmid # 60954) (17). This plasmid was PCR amplified with primers AT574 and AT575 using KAPA HIFI Mastermix (39). Primer sequences can be found in **Table S2**. The PCR product was purified using the Qiagen column purification kit. The two pieces were combined using the GeneArt Seamless PLUS Cloning and Assembly Kit at a ratio of 2:1 PCR product: plasmid digest. This generated a novel plasmid containing KRAB-dCAS9 with a P2A followed by mCherry under the tet operator. These were flanked with AAVS1 targeting homology arms as well as a puromycin resistance gene that is inserted in-frame in the AAVS1 gene. Complete plasmid sequencing was performed by Massachusetts General Hospital DNA Core by Next-Gen sequencing and automated sequence assembly. The clone used had a mutation in mCherry gene that resulted in a reduction in signal allowing for the utilization of mCherry as a selection marker for pUCM-CLYBL-*Ngn1/2* cassette integration sequentially after integration of the DOX-KRAB-dCAS9-puro cassette.

### Generation of iNeuron/CRISPRi cell line

Foreskin fibroblasts were transfected with the AAVS1-DOX-KRAB-dCAS9 plasmid and the 2 AAVS1 Talen plasmids (Gifts from Danwei Huangfu, Addgene plasmid # 59025 and 59026) using the Neon Transfection system (38). The settings were 3 pulses of 1,650 V for 10 ms (19). After two days, cells that integrated the target vector were selected by puromycin. PCR for integration was confirmed with one primer in the puromycin resistance gene and the other in the AAVS1 intronic region beyond the 500 bp homology arms of the plasmid resulting in a 1033 bp product (AT601 and AT602). The puromycin selected fibroblasts were then transfected with pUCM-CLYBL-*Ngn1/2* plasmid, a gift from Dr. Michael Ward and the 2 *CLYBL* targeting TALEN plasmids (pZT-C13-R1 and pZT-C13-L1, gifts from Jizhong Zou Addgene.org: #52638 and 52637, respectively) by the same method with the Neon Transfection System (40). The mCherry fluorescence was used for selection by fluorescence activated cell sorting. Correct integration of the cassette was confirmed by a similar PCR reaction (AT607 and AT608). Primer sequences can be found in **Table S2**. These fibroblasts were then reprogrammed identically to the protocol published previously (22).

### CRISPR gRNA virus production

The human CRISPRi-v2, genome-wide, top 5 sgRNAs/gene library was gift from Jonathan Weissman (Addgene #83969) (20). The library (100 µg) was electroporated into 50 µL of MegaX DH10B T1R Electrocompotent Cells (Invitrogen C640003) using a BioRad micropulser at 2KV. Dilutions onto bacterial plates were used to confirm adequate transformants for full library representation (∼6x10^9 transformants). Individual sgRNAs from the library were cloned by integrating synthesized oligos into the pCRISPRia-v2 plasmid (Addgene #84832) for the first test gRNA library. We switched and recloned in the pBA904 (Addgene #122238) which contains a capture sequence (cs1) in the loop of the sgRNA constant region used for single-cell RNA sequencing (41). Sequences were confirmed by Sanger sequencing. For each of the sub-libraries, these individual plasmids were mixed in equal molar amounts. Libraries and individual sgRNA lentiviruses were produced by the University of Michigan Vector Core.

### Quantitative Reverse Transcriptase PCR

Total RNA was isolated using the RNeasy Mini Kit (Qiagen). For cDNA synthesis, 1 µg of DNA was reverse transcribed with OligoDT using the SuperScript III kit (Invitrogen). Quantitative PCR was performed using the Power SYBR Green PCR Master Mix (Applied Biosystems, #4367659) with 10 ng of cDNA and 250 nM final primer concentration on a QuantStudio3 qPCR machine (Applied Biosystems). For each sample, actin primers were used as a loading control for a DDCT relative expression calculation. Primer sequences can be found in **Table S2**.

### Immunoblot analysis

Total protein from BFP expressing iNeurons was extracted using CelLytic™ M Cell Lysis Reagent (Sigma-Aldrich, #C2978) with PhosSTOP™ (Roche, #04906845001) and cOmplete™ ULTRA Protease Inhibitor Cocktail (Roche, #05892970001). Protein concentrations were determined by Qubit™ Protein BR Assay kit (Invitrogen, #A50668). Equal amounts of protein were denatured at 99°C for 5 minutes, separated by SDS-PAGE, and transferred onto NC membranes. The blots were blocked with 5% nonfat dry milk in Tris-buffered saline with 0.1% Tween 20 (TBS-T) for 1 hour, followed by incubation with primary antibodies at 4°C overnight (Antibodies and dilutions in **Table S3**). The blots were washed with TBS-T followed by incubation with IRDye 800 CW goat anti-rabbit or goat anti-mouse secondary antibodies (LI-COR Biosciences, 1:8000) for 1 hour. After additional washes with TBS-T, the immune complexes were detected by the Odyssey Imaging Systems (LI-COR Biosciences). Image Studio TM Lite Software (LI-COR Biosciences) was used to quantify the protein signals.

### iPSC culture and CRISPRi iNeuron differentiation

The KN-5 iPSC line was grown in 6-well plates coated with 1:200 Geltrex diluted in cold DMEM/F12 (1 mL/well) and incubated at 37°C for 30+ minutes. Cells were cultured in mTeSR1 with daily media changes. When cultures reached approximately 50% confluency, cells were passaged by incubating for 2 minutes in a sodium citrate solution and detached using a mini cell scrapper into media. Cells were triturated and replated at a 1:12 dilution.

To generate CRISPRi iNeurons, cell culture dishes were first coated with 1:50 Geltrex in DMEM/F12 for 30+ minutes at 37°C. The KN-5 line was washed with PBS and incubated in Accutase at 37 C for 8 minutes. The cells were resuspended in mTeSR1, centrifuged, and resuspended in mTeSR1 with 10 µM Y-27632. Cells were counted and added to the culture dishes at 1.5 x 10^5^ cells/cm^2^ in mTeSR1/Y-27632. The next day media was changed to mTeSR1 with 1 µg/mL dox. Immediately following, lentivirus containing the sgRNA or sgRNA library was added to the cells. For libraries, the MOI was targeted to be ∼ 0.3 to reduce multiplets. For individual sgRNA viruses, MOI between 0.6-0.9 was ideal for efficient use of cells and virus while reducing viral induced toxicity. For the 2 days following, the media was replaced with fresh mTeSr1 with dox.

### Live Fluorescence Activated Cell Sorting (FACS)

The third day following viral transduction, the cells are amenable to replating. Since the KN-5 line has a constitutive puromycin resistance gene, we were unable to utilize the puro selection that is built into the CRISPRia-V2 construct. Instead, we utilized the TagBFP fluorescent protein expressed on the construct as well to select for virally transduced cells. The cells were incubated with accutase for 8 minutes, resuspended in ice-cold 15% KOSR in PBS, counted, and centrifuged at 300 g for 5 minutes. The cells were resuspended at ∼ 5x10^6^ cells/mL. Cells were sorted for mCherry/TagBFP double-positive cells on a BD FACSAria II. Sorted cells were plated onto dishes coated with PEI and either Geltrex or laminin (1:100).

### pS6 intracellular staining and FACS

Cultures of day 7 iNeurons were incubated in accutase for 10 minutes. The cells were then triturated 3x with a glass Pasteur pipette, resuspended in cold PBS with 15% KOSR, and centrifuged at 1000 g. Cells were then resuspended in 1.5% paraformalydhyde in PBS at RT for 20 min. After centrifuging at 1000 xg for 5 minutes in a 5 mL FACS tube, the cells were resuspended in ice-cold 100% methanol at 0.5 mL/1x10^6^ cells. After vigorous vortexing, the cells were incubated on ice for at least 10 minutes and can be stored at 4 C for up to 1 week. On the day of FACS, the cells were centrifuged at 1000 g for 5 minutes followed by 2 washes with Biolegend Cell Staining Buffer. The cells were then incubated with pS6(S235/236)-Alexa647 conjugated primary antibody (1:500) for 30 min at RT in a volume of 100 µL/1x10^6^ cells in the staining buffer. The cells were then washed once in 5 mL of Cell Staining buffer and resuspended in the same buffer at 5x10^6^ cells/mL. The cells were sorted by Alexa647 fluorescence on the BD FACSAria II.

### Preparing gRNA PCR product for next-gen sequencing

Cells were pelleted by centrifugation at 1000 *g* for 5 minutes at 4°C. DNA was extracted following the Qiagen DNA FFPE extraction kit starting with the addition of 180 µL of ATL buffer and using a 30-minute reverse crosslinking step at 90°C. DNA was quantified and amplified using the KAPA HIFI kit with primers AT776 and AT881 with an annealing temperature of 65°C with 25-35 cycles based on DNA concentration. The PCR product was purified with Ampure beads and eluted in 25 µL of EB buffer.

### Calculating the log fold enrichment and rank order

The NGS data from the pS6-high and pS6-low populations was analyzed by 2 different methods. The first by gRNA in which gRNAs with fewer than 4 counts combined for the two populations were removed. The log fold enrichment score for each gRNA was calculated as log_2_([pS6-high +1]/[pS6-low +1]).

### In-cell western assay

On day 3 of dox treatment of CRISPRi iNeurons, cells were replated onto either PEI/Geltrex or PEI/laminin coated thin-bottom 96-well plates (Greiner µ-clear) at a density of 1x10^6^ cells/mL with 200 µL of media (3N+A+dox) added to each well. 100 µL of media was exchanged for fresh 3N+A media + 1 µg/mL dox each day. On day 7 the media was removed, and the cells were fixed with 4% paraformaldehyde in PBS at room temperature for 30 min. The cells were then washed 5 times for 10 min each with 200 µL of PBS with 0.1% Triton-X 100. The cells were then blocked for 1.5 hours in Odyssey Intercept Blocking Buffer (PBS). The cells were incubated with primary antibody at 1:400 in Odyssey blocking buffer with 50 µL in each well for 2.5 h. We performed no-primary control in 1 well each and no-cell control with primary antibody for 1 well each. Cells were washed 4 times x 5 minutes in PBS with 0.1% Tween-20. Secondary antibodies were incubated for 1 hour at RT with LiCor IRdye800 secondary antibody at 1:800 dilution in Odyssey blocking buffer along with 1:500 CellTag normalization dye. The cells were washed 5 times x 5 minutes in PBS with 0.1% Tween-20. All liquid was removed, and the plates were scanned on the Li-Cor Odyssey Imaging System. The In-Cell Western quantification analysis in the LiCor software was used to quantify each well for the 800 and 700 channels. The no-primary control values from the 800 channel were subtracted from all other wells with 800 channel values, and the no-cell values for the 700 channel. The 800 channel values for each well were divided by the 700 channel values. Finally, the values were normalized to the mean values of the No Target Control (NTC) wells (C145).

## Acknowledgments

We would like to acknowledge Louis Dang and Peter Crino for helpful conversations about the manuscript. We would also like to acknowledge the University of Michigan Flow Cytometry and Viral Vector Core Facilities for their important contributions to this research study. This work was funded by NIH/NINDS NS1162509 (AMT) and Citizens United for Research in Epilepsy (CURE) Innovator’s Award (JMP).

## Notes

### Competing Interest Statement

The authors have declared no competing interest.

