## Supplemental Figures and Tables for "Genome-wide CRISPRi Screen in Human iNeurons to Identify Novel Focal Cortical Dysplasia Genes"


Figure S1

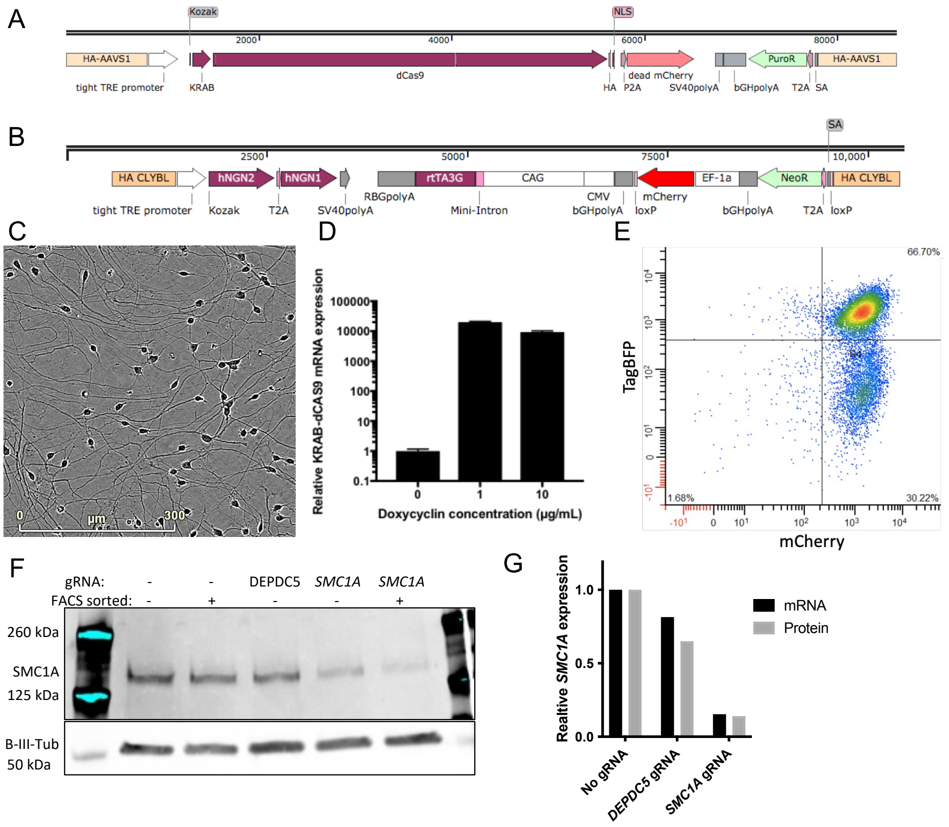


**Fig. S1.** Establishing the KN-5 CRIPSRi iPSC-derived iNeuron lines. (A) Schematic plasmid map depicting the TetO-promoter driven KRAB-dCas9 gene with puromycin resistance gene and AAVS1 targeting homology arms. (B) Schematic plasmid map depicting the TetO promoter-driven NGN1,2 along with the rtTA necessary for the Tet-On system, constitutive mCherry and Neomycin resistance genes for selection, and CLYBL targeting homology arms. (C) Phase micrograph of iNeurons on day 7 of doxycycline (dox) generated from KN-5 line. (D) Reverse transcriptase qPCR for KRAB-dCAS9 mRNA with 0, 1, or 10 μg/mL exposure to dox for 24 hours. (E) Flow cytometry data showing efficient TagBFP and mCherry signal to isolate healthy lentiviral transduced iNeurons. (F) Immunoblot for SMC1A protein and beta-III-tubulin loading control in day 7 iNeurons with either no gRNA transduction, *DEPDC5* gRNA or *SMC1A* gRNA, and with (+) or without (-) FACS for TagBFP/mCherry double-positive iNeurons. (G) qRT-PCR for *SMC1A* transcript (black bars) and normalized quantification of SMC1A protein (gray bars) from the immunoblot in Figure S1F.

Figure S2

A

| | gRNA | GAACTCGTCATCCACAAGAA |
|---|---|---|
| Line Con2 | WT/WT | ACAAACTCGTCATCCACAAGAAGGGCTTT |
| Line D7 | -20/-20bp | ACAAA--------------------CTTT |

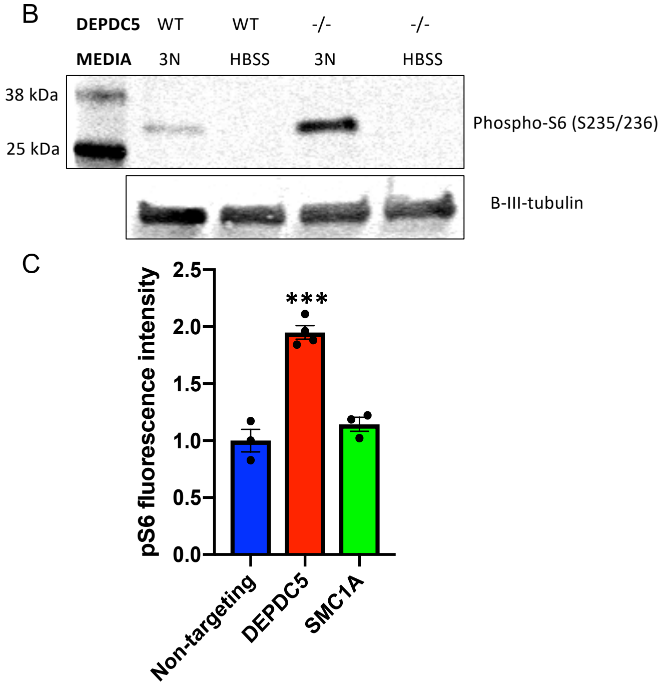


**Fig. S2.** Homozygous *DEPDC5* frameshift mutation causes elevated pS6 (S235/S236) levels in iNeurons. (A) gRNA used for CRISPR editing and genotype data for two lines that underwent CRISPR-based mutagenesis of *DEPDC5* and simultaneous reprogramming. These lines were also incorporated with the iNeuron CLYBL insertion vector via TALENs and mCherry selection. (B) Immunoblot for pS6(S235/S236) from the two lines differentiated into iNeurons by dox exposure for 7 days. b-III-tubulin was used as a loading control. (C) In-cell western assay for pS6 in iNeurons transduced with NTC, *DEPDC5,* or *SMC1A* gRNAs. Each dot is data from one well. Statistical analysis performed by one-way ANOVA with Dunnet's multiple comparison test. P *** < 0.001. Error bars are mean +/- STD.

Figure S3

A

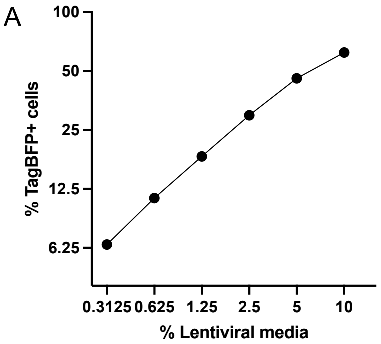


B

| | Before FACS | | After FACS | |
|---|---|---|---|---|
| MOI | % Singlets | % Multiplets | % Singlets | % Multiplets |
| 0.5 | 25 | 25 | 50 | 50 |
| 0.4 | 24 | 16 | 60 | 40 |
| 0.3 | 21 | 9 | 70 | 30 |
| 0.2 | 16 | 4 | 80 | 20 |
| 0.1 | 9 | 1 | 90 | 10 |

**Figure S3.** Titrating the gRNA containing lentivirus by TagBFP FACS and MOI modeling. (A) iNeurons were transduced with a log-2 dilution series of lentivirus containing media ranging from 0.3125% up to 10% of the culture media. The percentage of TagBFP+ cells was determined by FACS. Both axes are plotted in log-2 format. (B) Table modeling the number of singlet and multiplet gRNA transduced cells based on the assumption that the probability of double transduction is the $MOI^2$ and so on. The ideal MOI was targeted to be between 0.1-0.3 to reduce the number of multiplets during library screens.

Figure S4

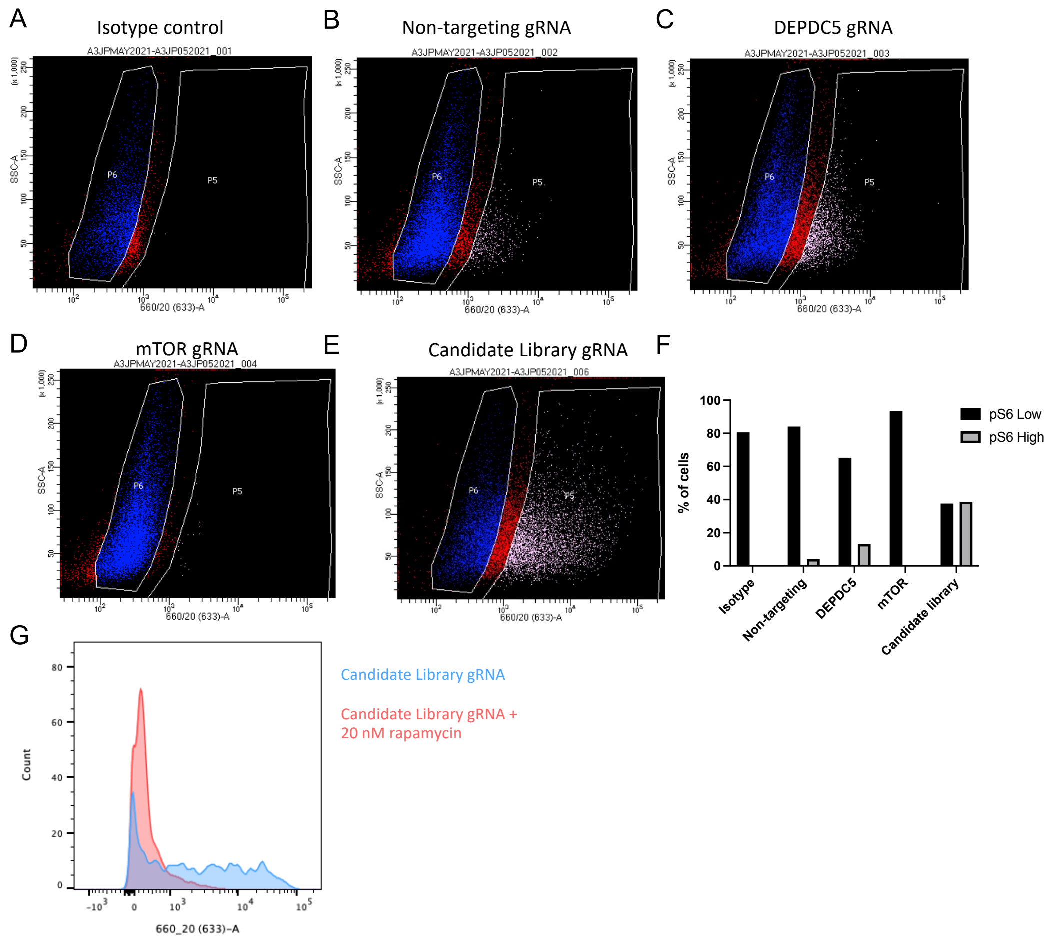


**Figure S4.** Representative FACS plots sorted for pS6-high and pS6-low iNeuron populations under different conditions. (A-E) iNeuron FACS plots with side-scatter area in the y-axis and pS6-Alexa-647 fluorescent intensity in the x-axis. The sorted pS6-high population is in pink, the sorted pS6-low population is in blue, and the discarded cells are in red. (A) Cells were incubated with an isotype control antibody conjugated with Alexa-647 to see background signal in this channel due to non-specific antibody binding. (B) iNeurons transduced with a non-targeting control (NTC) gRNA and incubated with pS6-Alexa647. (C) iNeurons transduced with a *DEPDC5* gRNA and incubated with pS6-Alexa647. (D) iNeurons transduced with a mTOR gRNA and incubated with pS6-Alexa647. (E) iNeurons transduced with the candidate gRNA library and incubated with pS6-Alexa647. (F) Graph of the percentage of total cells in the pS6-high and pS6-low sorted groups for each condition in (A-E). (G) FACS histogram of iNeurons transduced with the candidate gRNA library and treated for 2 hours with either 20 nM rapamycin or vehicle.

Figure S5

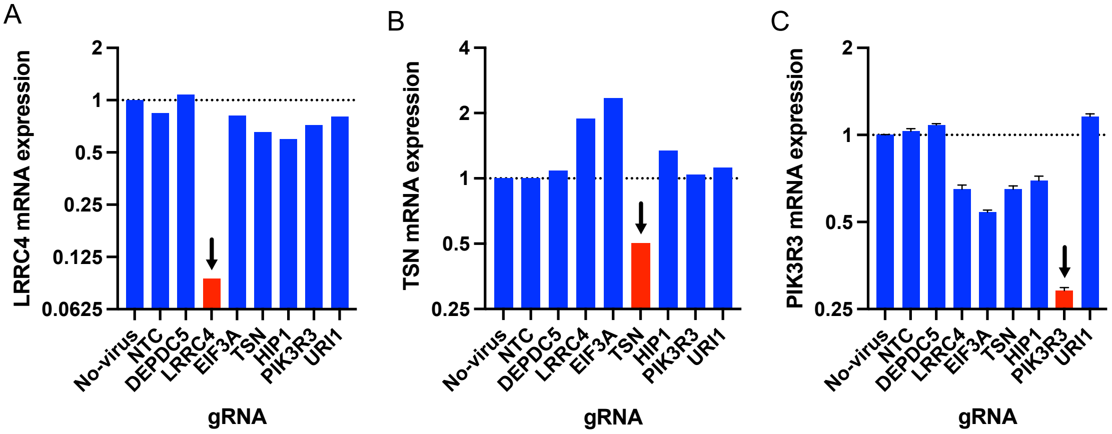


**Figure S5.** Reduced mRNA transcript levels after lentiviral CRISPRi on-target gRNA expression for a subset of gene hits from the whole-library gRNA screen. (A) Quantitative RT-PCR for *LRRC4* mRNA in day 7 iNeurons transduced with the various gRNAs shown on the x-axis. Reduced transcript abundance is only seen for the *LRRC4* targeting gRNA. (B) Quantitative RT-PCR for *TSN* mRNA in day 7 iNeurons transduced with the various gRNAs on the x-axis. Reduced transcript abundance is only seen for the *TSN* targeting gRNA. (C) Quantitative RT-PCR for *PIK3R3* mRNA in day 7 iNeurons transduced with the various gRNAs on the x-axis. Greatest reduction in transcript abundance is seen for the *PIK3R3* targeting gRNA. *N* = 1 for each condition in all three graphs.

Figure S6

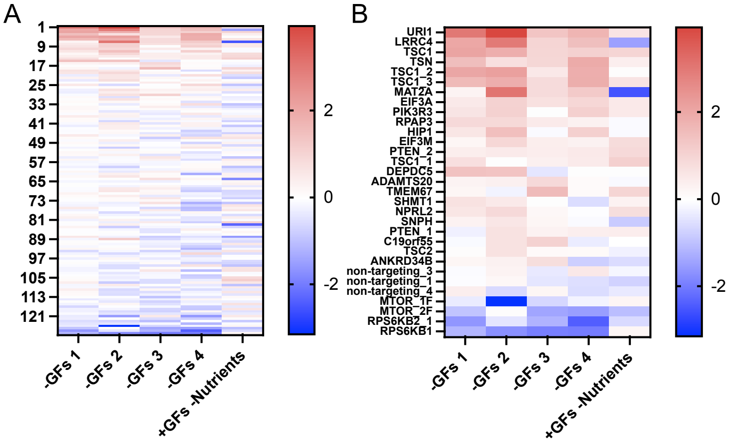


**Figure S6.** Heatmap for log2 fold enrichment for gRNA candidate library for 5 experiments. (A) Log2 fold enrichment heatmap for all 130 genes in the gRNA candidate library for the 4 independent experiments with growth factor withdrawal and one experiment with nutrient withdrawal. (B) Log2 fold enrichment heatmap for all significant candidate genes, known FCD genes, and NTCs from the gRNA candidate library for the 4 independent experiments with growth factor withdrawal and one experiment with nutrient withdrawal.

| Table S1. Test Library | | |
|---|---|---|
| gRNA # | Gene ID | gRNA sequence |
| C97 | STRADA | CAGTTTTACTGCCGCGCCG |
| C99 | STRADA | CGACCGTCTTGGCTCCGCC |
| C101 | DEPDC5 | CCAGATACCCCGCCTACGC |
| C103 | DEPDC5 | CTCCGGTGGCTCCGCCCCG |
| C105 | TSC1 | TCGTGAAAGGGCCAAGGCC |
| C107 | TSC1 | TCAGCTGTTTACCTCACAG |
| C109 | TSC2 | CGTGCGGCCCTGCGTTCCC |
| C111 | TSC2 | CTGCCCCCTTTCGCCCCAA |
| C113 | PTEN | CTCGGAAGCTGCAGCCATG |
| C115 | PTEN | CATCTCTCATCTCCCTCG* |
| C117 | RPS5KB1 | CCATCACCGGCTGCTTAGG |
| C119 | RPS5KB1 | GTGATGGCGGCAGCGGCTG |
| C123 | RPS5KB2 | GGACTGTCAGTCAGTGCGC |
| C125 | MTOR | GCTGTCCTCTAAGCCGGGA |
| C127 | MTOR | CCCGCCTTCCCCGCTGTCC |
| C129 | AKT3 | tcgcttgccctcccGCCGG |
| C131 | AKT3 | ttgcaggtaacagccaccg |
| C133 | HPRT1 | GCTCCGTTATGGCGACCCG |
| C135 | HPRT1 | TTATGGCGACCCGCAGCCC |
| C137 | CLYBL | CCCTCCGCAGCAGACGTAG |
| C139 | CLYBL | TCCGCAGCAGACGTAGCGC |
| C141 | AAVS1 | GGGCCACTAGGGACAGGAT |
| C143 | AAVS1 | ATCCTGTCCCTAGTGGCCC |
| C145 | non-targeting | TGATTCGCGCCCCATGCAG |
| C149 | non-targeting | CAACGGCGGAGGTTGCACA |
| C153 | non-targeting | CTCAATCCCTCTACCTGAC |
| C155 | non-targeting | CTGGAGTACGATCGTCCTC |
| *The NGS reads for this sequence have an A where the designed gRNA and human genome have a G. | | |

**Table S1. Test gRNA library.** A small lentiviral library containing 27 gRNAs. The green gRNAs target known negative mTOR regulator genes, red gRNAs target kinases that should increase pS6, and black gRNAs should not affect pS6 levels.

| Table S2. Primer sequences | | |
|---|---|---|
| **Primer Name** | **Description** | **Sequence** |
| AT103 | *Actin* F1 | CTGTGGCATCCACGAAACTA |
| AT104 | *Actin* R1 | AGCACTGTGTTGGCGTACAG |
| AT351 | *WPRE* F1 | CTTCCCGTATGGCTTTCATT |
| AT352 | *WPRE* R1 | GGGCCACAACTCCTCATAAA |
| AT886 | Myco F SC1 | TGCACCATCTGTCATTCTGTT |
| AT887 | Myco R SC2 | GGGAGCAAACAGGATTAGATA |
| AT589 | *DEPDC5* F1 | GCACACAGGTTTGGGTTTG |
| AT590 | *DEPDC5* R1 | AGTAGGGCAGCTGCAGAAAC |
| AT595 | *TSC1* F1 | ACAGGCAGCTGTTGGTTCTT |
| AT596 | *TSC1* R1 | TAGGCGGCTTTCATCATTTC |
| AT591 | *HPRT1* F1 | GACCAGTCAACAGGGGACAT |
| AT592 | *HPRT1* R1 | CTGCATTGTTTTGCCAGTGT |
| AT561 | *SMC1A* F1 | TCAGAGCCGAGAGAGGGAAA |
| AT562 | *SMC1A* R1 | TGAATGCCCAAGCGAGTCTT |
| AT605 | *dCAS9* F1 | CTTGCTGGGCACCTTGTACT |
| AT606 | *dCAS9* R1 | GAGGGTCGATGGACAAGAAG |
| AT492 | *DEPDC5 sequencing* F1 | GCAGGGAGGCAAGATGACTT |
| AT493 | *DEPDC5 sequencing* R1 | TGCACACACCAGAACCTCAG |
| AT599 | *gRNA lentiviral titering* F1 | CAGCACAAAAGGAAACTCACC |
| AT600 | *gRNA lentiviral titering* R1 | CGACTCGGTGCCACTTTT |
| AT599 | *gRNA amplification for CRISPRia-V2 F* | CAGCACAAAAGGAAACTCACC |
| AT600 | *gRNA amplification for CRISPRia-V2 R* | CGACTCGGTGCCACTTTT |
| AT776 | *gRNA amplification for CRISPRia-V2 F with Illumina adapter* | TCGTCGGCAGCGTCAGATGTGTATAAGAGACAGCAGCACAAAAGGAAACTCACC |
| AT777 | *gRNA amplification for CRISPRia-V2 R with Illumina adapter* | GTCTCGTGGGCTCGGAGATGTGTATAAGAGACAGCGACTCGGTGCCACTTTT |
| AT880 | *gRNA amplification for pBA904 R* | CTTAAAGCggccAAGTTGAT |
| AT881 | *gRNA amplification for pBA904 R with Illumina adapter* | GTCTCGTGGGCTCGGAGATGTGTATAAGAGACAGCTTAAAGCggccAAGTTGAT |
| AT601 | *AAVS1 insertion boundary* | gtgggcttgtactcggtcat |
| AT602 | *AAVS1 insertion boundary* | CTGCCGTCTCTCTCCTGAGT |
| AT607 | *CLYBL insertion boundary* | gtaaaccactgtggggtgga |
| AT608 | *CLYBL insertion boundary* | AGCAAAAGACCCGACTCAGA |
| AT554 | *CLYBL flanking insertion site* | CAAACATGGCTCAGTTGTGAA |
| AT555 | *CLYBL flanking insertion site* | TTGGTGGTGGTCACAGTCAT |
| AT575 | KRABdCas_F1 | CAAGAAAGCTGGGTCGTACTTGTACAGCTCGTCC |
| AT574 | KRABdCas_R1 | TTTCAGGTTGGACCGGCGCGTCTGCAGGATA |
| AT576 | SV40pA-R Sequencing Primer | GAAATTTGTGATGCTATTGC |
| AT577 | attB1 Sequencing Primer | ACAAGTTTGTACAAAAAAGCAGGCT |

**Table S2. Primer sequences.** This list contains all qRT-PCR, PCR, and sequencing primers used in this study.

| Table S3. Antibodies and dilutions | | | | |
|---|---|---|---|---|
| **Antigen** | **Species** | **Dilution** | **Vendor** | **Catalog#** |
| **pS6(S235/236)** | Rabbit | 1:2000 | Cell Signaling Technology | 4858 |
| **pS6 (Ser235/236) Alexa Fluor® 647** | Rabbit | 1:400 | Cell Signaling Technology | 4851 |
| **S6** | Rabbit | 1:1000 | Cell Signaling Technology | 2217 |
| **SMC1** | Rabbit | 1:2000 | Bethyl | A300-055A |
| **EIF3A** | Rabbit | 1:1000 | Cell Signaling Technology | 3411 |
| **URI/RMP** | Rabbit | 1:1000 | Thermo | 11277-1-AP |
| **Raptor** | Rabbit | 1:1000 | Cell Signaling Technology | 2280 |
| **HIP1** | Rabbit | 1:1000 | Cell Signaling Technology | 90830 |
| **Tubulin β3** | Mouse | 1:10000 | Biolegend | 801201 |
| **P-p70S6k(T389** | Rabbit | 1:1000 | Cell Signaling Technology | 9205 |
| **pAKT (S473)** | Rabbit | 1:1000 | Cell Signaling Technology | 9271 |
| **pMTOR(S2448)** | Rabbit | 1:1000 | Cell Signaling | 5536 |
| **pERK (T202/Y204)** | Rabbit | 1:1000 | PhosphoSolutions | 160-2024 |
| **p-4E-BP1** | Rabbit | 1:1000 | Cell Signaling Technology | 2855 |

**Table S3. Antibody dilutions.** This table contains the name, host species, dilution, company, and catalog information for all antibodies used in this study.

**Dataset S1 (separate file).** This file contains the data for the CRISPR screen experiments. The first tab contains the rank order for the 3 independent whole genome experiments for each gene and an average rank order. Tab 2 contains the raw NGS reads for each gRNA in each of the 3 independent whole genome screen experiments along with the $log_2$ fold change (LFC), average LFC, and rank order. Each gRNA sequence is also listed. Tab 3 contains normalized NGS reads for the 3 candidate library screens. The fold change for each gRNA for each experiment is listed along with the average LFC and rank order. Each gRNA sequence is also listed.
